# Urine proteomics as a non-invasive approach to monitor exertional rhabdomyolysis during military training

**DOI:** 10.1101/2021.12.10.471761

**Authors:** Andréia Carneiro, Janaína Macedo-da-Silva, Verônica Feijoli Santiago, Gilberto Santos Oliveira, Thiago Guimarães, Clarissa Ferolla Mendonça, Jéssica Laís de Oliveira Branquinho, Cintia Verdan Lucena, Juliana Batista Osório, Eduardo Pernambuco, Josino Costa Moreira, João Bosco Pesquero, Marcos Dias Pereira, Giuseppe Palmisano

**Author notes:** Equal contributions (Andréia Carneiro, Janaína Macedo-da-Silva). Authors to whom correspondence should be addressed, Corresponding author: Prof. Giuseppe Palmisano; Postal address: Av. Prof. Lineu Prestes, 1374, CEP: 05508-900 - São Paulo – SP – Brazil. Prof. Marcos D. Pereira; Postal address: Avenida Athos da Silveira Ramos, 149, Bloco A, 5° andar, Lab. 549-C, CEP: 21.941-909, Cidade Universitária, Rio de Janeiro, RJ, Brasil; Rel. Andréia Carneiro; Postal address: Rua Primeiro de Março, 118, 17° andar, Centro, CEP: 20.010-000, Rio de Janeiro, RJ, Brasil.

## Abstract

Exertional rhabdomyolysis (ERM), a condition often associated with strenuous exercise, a common practice in the military activities, can be defined as the process of injury and rupture of muscle cell membranes, with leakage of its components into the blood stream. Creatine kinase (CK) has been extensively used for ERM diagnosis, albeit several studies reported the discrepancy between CK levels and clinical signs or symptoms. In this study, we analyzed the biochemical profile of the blood, and the urinary proteome of ten marine soldiers in a special training course. The samples were collected in two periods, M1 and M2, which correspond to the lowest and highest CK levels during training, respectively. Quantitative urinary proteome profile of M1 and M2 was determined showing changes with highest significance in immune system and cell adhesion-related pathways after strenuous physical exercise. Changes in the abundance of several proteins was observed in individuals carrying genetic polymorphisms related to greater risk for muscle damage. Remarkably, we identified a panel of proteins (CTSH, PIK3IP1, DEFB1, ITGB1, BCAN, and TNFRSF10C) that present high correlation with three classical blood biochemical markers of ERM and *AGT* MET235Thr and *ACE* I/D polymorphisms. These proteins represent potential urine markers of muscle damage due to intense physical conditions such as military training activities.

## Introduction

The practice of physical exercise has been related to several health benefits; however, if performed vigorously, prolonged and with high loads, it may cause serious damage to the muscle tissue (Rhodes et al., 2017, Peluso and Andrade, 2005, Warburton et al., 2006). In this context, professional, amateur, or recreational athletes are often affected by metabolic, immunological, neurological, endocrine, cardiovascular, musculoskeletal, and renal changes due to strenuous physical effort (Fernandez et al., 2018, Ciolac and Guimarães, 2004, Johansen, 2005). Strenuous physical exercise has been associated with damage to the striated muscle with the subsequent release of its cellular constituents (e.g. electrolytes, myofibrillar, cytoskeletal, and membrane proteins) into the blood stream (Owens et al., 2019, Howatson and van Someren, 2008). The etiology of this rupture is complex and diverse, which includes physical effort, muscle hypoxia, genetic susceptibility, infections, changes in body temperature, and drug use (Owens et al., 2019).

Rhabdomyolysis (RM), can be defined as the process of injury and rupture of muscle cell membranes, with leakage of its components into the blood stream (Torres et al., 2015, Chavez et al., 2016, Efstratiadis et al., 2007). The appearance of muscle pain, changes in sensitivity, muscle weakness, dark brown urine, myalgia, and edema of body segments are classic symptoms for RM (Torres et al., 2015, Efstratiadis et al., 2007). The main consequences of RM may be associated with acute renal failure, electrolytic disorder, cardiac arrhythmia, compartment syndrome, and eventually death (Sousa et al., 2013, Chen et al., 2013, Caldeira et al., 2001). The incidence of RM by physical exertion is approximately 30 for every 100,000 patients per year (Danielak et al., 2018, Eichner, 2020). The sport that stands out the most in relation to the number of investigations on exertional RM (ERM) is running (Brinley et al., 2018). However, the literature also points to the development of ERM in farmers and military personnel, especially those exposed to exertional work activities and thermal stress (Peraza et al., 2012, Moyce et al., 2017). Data from 2019 revealed that amongst US armed service members the occurrence of ERM is 38.9 per 100,000 people/year. In addition, the rate in the Marine Corps subgroup is 91.9 per 100,000 people/year, with the youngest and the least time on duty, those of African descent, and those with less education being most affected (Carneiro et al., 2021).

When ERM is suspected, physical activity should be stopped and the patient subjected to blood and urine tests to determine the levels of traditional markers [e.g. myoglobin (MB), creatine kinase (CK), aspartate transaminase (AST), aldolase, alanine transaminase (ALT), and lactate dehydrogenase (LDH)] used for diagnosis (Cervellin et al., 2010, Keltz et al., 2013, Nance and Mammen, 2015). Traditionally, the CK has been used as the main diagnostic biomarker for RM (Nance and Mammen, 2015). Once the identified levels of CK, associated or not with other markers, are higher than indicated under normal conditions, the patient must be taken to the health system (Nance and Mammen, 2015, Landau et al., 2012). However, there are several reports that describe the discrepancy between high levels of CK and clinical signs or symptoms of this disorder, while there are people with low levels of this enzyme with rhabdomyolysis (Luckoor et al., 2017). It is suspected that the individual genetic background may explain this variation. Therefore, levels considered normal should be adapted according to the individual’s genotype (Scalco et al., 2015). The genetic variants for angiotensinogen (*AGT*) and angiotensin conversion enzyme (*ACE*) were correlated to muscle damage and skeletal muscle efficiency. Several hereditary metabolic disorders and some genetic variants, including genes that code for *AGT* and *ACE*, are now recognized as contributing to an increased risk of rhabdomyolysis (Landau et al., 2012), being associated with increase in CK. Studies have linked the I-allele *ACE* and the increase in CK after intense physical activity with a higher risk of developing ERM (Yamin et al., 2007).

Recently, it has been shown that the renin-angiotensin system plays an important role in the loss of skeletal muscle mass. *AGT* is cleaved by renin into angiotensin I, which in turn is converted into vasoconstrictor angiotensin II by *ACE*, causing the degradation of bradykinin (vasodilator) (Sierra et al., 2019).

It is estimated that up to a third of cases of ERM evolve to acute impairment of renal function (Holt and Moore, 2001), which is one of the most severe outcomes of this disease. Renal damage is associated with increased levels of MB in the bloodstream, which results in renal vasoconstriction, the formation of intratubular cylinders, and direct toxicity to renal tubular cells (Kasaoka et al., 2010). The evaluation of potential markers of kidney and muscle damage, as well as the discussion about new diagnostic possibilities, seems to be highly relevant and may be useful for understanding the impact of ERM on tissues and/or organs functions (Ulusoy et al., 2013). In this context, the aim of this study was to identify potential urinary biomarkers to monitor ERM in military personal after vigorous and prolonged exercise during two military missions. Analysis of blood biochemical parameters, polymorphisms screening, and urinary proteome profiling identified differently regulated proteins associated with strenuous military physical training.

## Materials and Methods

### 1. Study design and exercise training program

A longitudinal/prospective study was performed including Brazilian marine soldiers enrolled in a special training program from April to December 2018 involving four phases. Clinical and demographic parameters of the volunteers, male and asymptomatic, were measured. The mean age (29,0 ± 2,7 years) and anthropometric measures total body mass (kg): 79,9 ± 7,1; height (cm): 177,2 ± 6,3; body mass index (kg·m-2): 25,3 ± 1,8; Body Fat Percentage (%): 10,4 ± 2,9: Lean body mass (kg): 71,6 ± 5,8; Peak VO2 (mL·kg-1·min-1): 52,3 ± 2,0 indicates homogeneity of samples. The samples of ten (10) volunteers were collected in two different moments of the military training, M1 and M2 (**Supplementary Figure 1**). Based on blood CK levels, M1 and M2 were defined as missions with low and high skeletal muscle damage, respectively.

This project was approved by the Research Ethics Committee (CEP) number 2.219.303 of Hospital Naval Marcilio Dias, Rio de Janeiro, Brazil. The militaries were individually invited to participate in the study and signed the written informed consent form. The current use of statins was an exclusion criterion.

### 2. Biofluids collection

Blood and urine samples were collected during the M1 and M2 missions, transported in a cold thermal box (4°C) and later stored at −80°C. Three blood samples per participant were collected from the antecubital arm using a disposable 20-gauge needle equipped with a Vacutainer tube holder (Becton Dickinson, Franklin Lakes, NJ). Two serum samples were collected (16 mL) and kept in Vacutainer tubes containing SST-Gel and in Vacutainer tubes containing EDTA anticoagulant. The urine samples (50-90 mL) were collected in a numbered and sealed plastic bottle and added protease inhibitor cocktail in the aliquots that were stored in the freezer −80°C.

### 3. Hematological and biochemical analyses

The following hematological biomarkers were analyzed using Symex XT-2000i: leucocytes and biochemical biomarkers: creatine kinase (CK) and its isoforms (CKMB), phosphorus (P), aspartate transaminase (AST). Lactate dehydrogenase (LDH) was measured using a Vitros 4600 Chemical System (Ortho-Clinical Diagnostics, Johnson and Johnson, Rochester, NY).

### 4. Sample preparation for proteomics analysis

Urine samples (10 mL) were centrifuged for 15 minutes at 2,000xg and 4°C. Then, the samples were filtered with Sterile Syringe filter (0.22 μM) and concentrated with Millipore Amicon Ultra-15 10K (Merck Millipore) according to the manufacturer’s instruction. The retentate was collected and the filter was washed with 200 μl of 4 M Urea and combined. This protein solution was quantified using the Qubit Protein Assay Kit (Invitrogen) platform. A volume corresponding to 100 μg was retrieved for each sample and 50 mM triethylammonium bicarbonate (final concentration) was added. Proteins were reduced with 10 mM dithiothreitol (DTT) for 40 minutes at 30 °C and alkylated with 40 mM iodoacetamide (IAA) for 30 minutes at room temperature in the dark. Samples were diluted to 0.8 M Urea (final concentration) and digested with trypsin (Promega) (1:50 w/w) for 16 hours at 30°C. After digestion, all reactions were acidified with 1% (v/v) trifluoroacetic acid, and tryptic peptides were purified using OASIS (HLB-SPE) cartridge. The samples were dried and suspended in 0.1% formic acid (FA).

### 5. Mass spectrometry analysis

Tryptic peptides (1 μg) were analyzed with an LTQ-Orbitrap Velos ETD (Thermo Fisher Scientific) coupled with Easy NanoLC II (Thermo Scientific). The peptides were separated on a ReproSil-Pur C18-AQ C18 reversed-phase pre-column (4cm × 100μm inner diameter, 5μm particles) and subsequently eluted onto a 20 cm 75 inner diameter analytical column containing ReproSil-Pur C18-AQ 3 μm particles over 105 minutes using a linear gradient 2-30 % followed by 20 minutes of 30-45% of mobile phase B (100% ACN; 0,1% formic acid). To acquire MS data, the data-dependent acquisition (DDA) mode operated by the XCalibur software (Thermo Fisher) was used. Survey scans (350–1,500 m/z) were acquired in the Orbitrap system with a resolution of 60,000 at *m/z*. The selection of top 20 precursors followed by fragmentation using collisional induced dissociation (CID) method was employed using normalized collision energy of 35. The MS/MS settings included: min. signal required of 5000, isolation width of 2.00, activation Q of 0.250 and activation time of 10 ms. A technical duplicate for each sample was run.

### 6. Data processing

The tandem mass spectrum data was searched using the Andromeda and MaxQuant v.1.5.5.135 (Tyanova et al., 2016) search algorithm. The *H. Sapiens* Swiss-Prot (Uniprot, 20,359 entries, reviewed) database was downloaded on September 20th, 2020. The search parameters were set to allow two missed tryptic cleavages; the oxidation of methionine and carbamidomethylation of cysteine was set as variable and fixed modifications, respectively. Other search criteria included: mass tolerance of 20 ppm (precursor ions) and 0.5 Da (fragment ions), and minimal peptide length of 7 amino acids. A false discovery rate (FDR) of 1% was applied at peptide and protein level. The “match between run” option was enabled. Protein quantification was performed using label-free quantification (LFQ) by integrating the extracted ion chromatogram (XIC) area.

### 7. Genotype determination

An aliquot of 125 μl was pippeted from EDTA tube on FTA cards (WhatmanTM – GE Healthcare ® - Cat. No. WB12 0206; Little Chalfont, United Kingdom). gDNA from blood were extracted using the Quelex 20% and Proteinase K (10 mg/mL) protocol and was quantified using a NanoDrop One (Thermo Fisher Scientific). Polymorphisms genotyping for the identification of the *AGT* Met235Thr were performed by qPCR allelic discrimination using commercial fluorescence-based TaqMan SNP Genotyping Assays (rs.699, Applied Biosystems, Foster City, CA, USA). Amplification and analyzes were performed in PCR thermocycler, QuantStudio™ 5 Real-Time PCR System (Thermo Fisher Scientif). Polymorphisms genotyping for identification of the *ACE* insertion (I) or deletion (D) variants were screened by a polymerase chain reaction (PCR) using a sense primer (5’-GAT GTG GCC ATC ACA TTC GTC AGA T -3’) and an antisense primer (5’-CTG GAG ACC ACT CCC CAT CCT TTC T− -3’). The PCR product resulted in a 490 bp (I) and 190 bp (D) fragment analyzed on a 2% agarose gel stained with SYBR® Safe DNA gel stain (Invitrogen).

### 8. Statistical analysis

All datasets were tested for normal distribution in order to apply parametric tests. For the proteomic data, the protein expression data were processed using the Perseus computational platform v.1.6.14.0 (http://www.coxdocs.org/doku.php?id=perseus:start). LFQ data were log2 transformed, protein reverse, contaminants and only by site were removed. Imputation was performed by replace missing values from normal distribution with a width of 0.3 and down shift of 1.8. Differentially regulated proteins between M1 and M2 were determined using paired t-test of LFQ values and correcting for Benjamini-Hochberg based FDR (FDR < 0.05). The comparison of differentially regulated proteins between the investigated genotypes was determined by applying a student t-test. The correlation between biochemical markers and regulated proteins was determined by applying the Spearman correlation test. Graphpad Prism 5.01 was used to perform statistical test analysis and considered significant with a p≤0.05.

### 9. Bioinformatics analysis

The analysis of gene ontology (GO) was determined by the ClueGO app. A q-value threshold of 0.05 was used, corrected by the Benjamini-Hochberg method (Bindea et al., 2009). The enriched pathways were determined by applying a Gene Enrichment Analysis (GSEA) using the Reactome and Wikipathways platform as reference data (Subramanian et al., 2005). Complementary pathways analyzes were performed using the g: profiler tool (Reimand et al., 2007), applying a 0.05 adjusted p-value significance threshold (Benjamini-Hochberg method). ROC curves were created using the pROC package (Robin et al., 2011), available from Bioconductor. Complementary analyzes were performed using Perseus, Graphpad prism v.8, and, RStudio software.

## Results

### Differently regulated proteins related to strenuous exercise in military activity

The workflow adopted for blood and urine samples analysis is shown in **Figure 1A**. The applied LFQ approach identified and quantified 548 urinary proteins. A total of 226 proteins were differentially regulated between M1 and M2, with 120 upregulated and 106 downregulated (**Figure 1B**). Principal component analysis (PCA) showed a clear separation between the two groups, revealing distinct proteomic profiles (**Figure 1C**). SOD1, FABP3, CA1, and CTSA proteins showed the greatest increase in fold change amongst upregulated proteins (**Figure 1D**). On the other hand, the downregulated proteins MSMB, SPIT2, CD248, and HMCN1 in M2 presented the greatest reduction in fold change (**Supplementary File 1**). Bioinformatics analysis was carried out to determine the tissue expression of differentially regulated proteins (**Figure 1F**). It is possible to observe a greater variety of proteins associated to different tissues, including proteins identified in the plasma, urine, liver, and skeletal muscle. Moreover, diseases associated with differentially regulated proteins were also determined (**Figure 1G**), and it was possible to identify several proteins related to the immune system, muscle weakness, motor delay, muscle spasm, skeletal muscle atrophy, amongst others.

**Figure 1.**
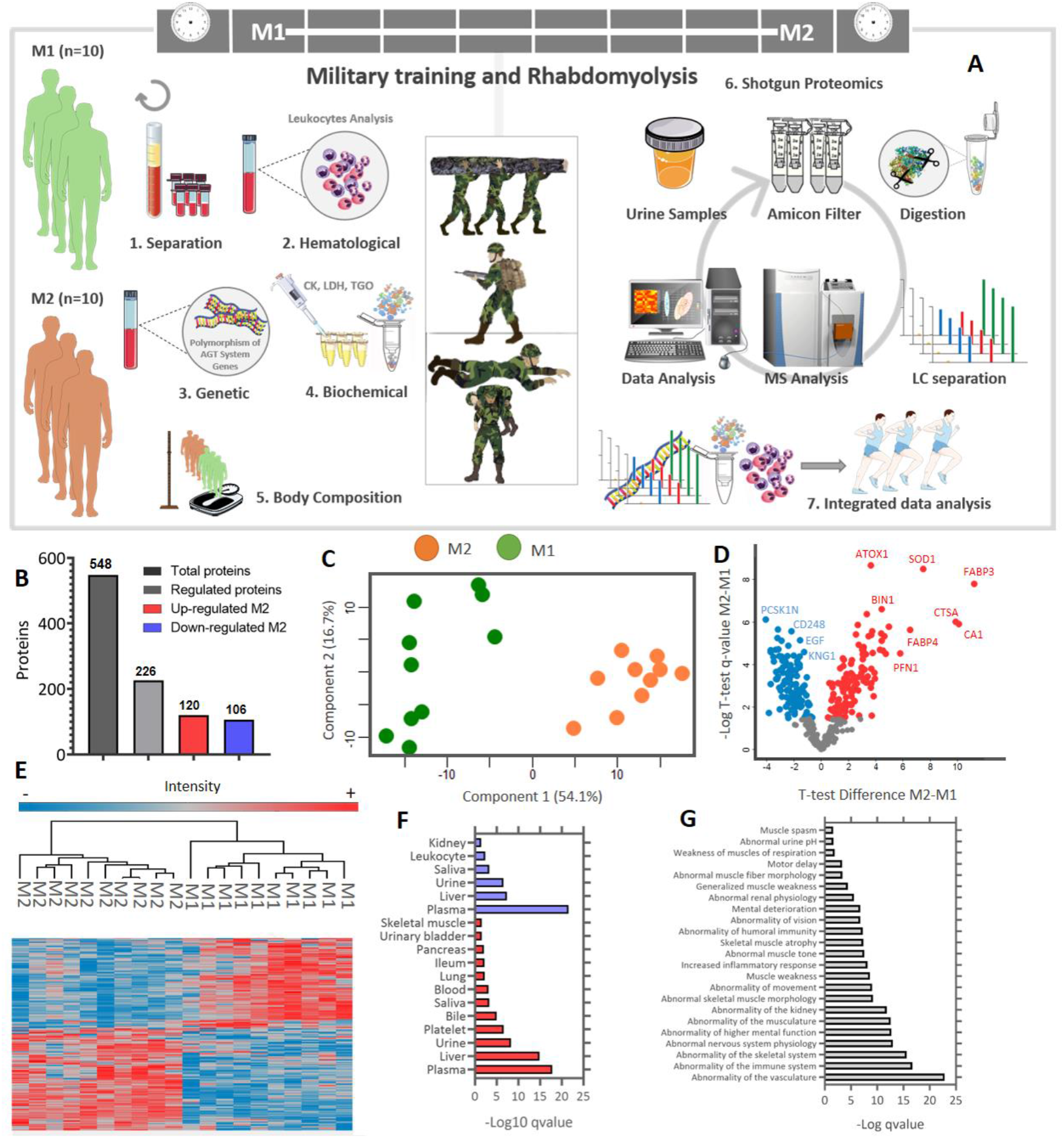
Experimental workflow to identify and validate differentially regulated urinary proteins in 10 soldiers during the M1 (low-physical stress) and M2 (high-physical stress) missions. Blood and urine samples were collected to perform biochemical, genetic, and proteomic analyses. The M1 and M2 groups were submitted to genetic, hematological, biochemical, and proteomic analysis (A). The samples were analyzed in an LTQ Orbitrap Velos system and the data obtained were mapped by the MaxQuant software, using the reviewed *Homo sapiens* database. Differentially regulated proteins were determined by applying a paired student t-test, with Benjamini-Hochberg correction (q-value ≤ 0.05). A filter to select only proteins with eight valid values in at least one group was previously applied (B). Principal component analysis of the total identified and quantified proteins in M1 and M2 (C). Volcano plot of differentially regulated proteins between M1 and M2. The foldchange is represented by the log2 ratio between the protein intensities M1 and M2; negative values indicate greater abundance in the M1 group compared to the M2 group (D). Heatmap of differentially regulated proteins. The data were normalized using the z-score function. Positive values indicate that the data is above average and when it is negative it means that the data is below average. The red and blue colors show positively regulated and negatively regulated proteins, respectively. (E). Tissue expression of upregulated (red) and downregulated (blue) proteins. Only terms with q-value ≤ 0.05 were represented (Benjamini-Hochberg) (F). Diseases associated with differently regulated proteins. Only terms with q-value ≤ 0.05 were represented (Benjamini-Hochberg) (G).

### Proteomics reveals dysregulation of the immune system and proteins related to the extracellular matrix (ECM)

The differentially regulated proteins showed significant enrichment for the following Uniprot keywords: Immunoglobulin domain, cell adhesion, pyrrolidone carboxylic acid, immunity, extracellular matrix, protease, GPI-anchor, serine protease inhibitor, adaptive immunity, calcium, lysosome, hydrolase (**Figure 2A**). The terms with the highest number of proteins were hydrolase, immunoglobulin domain, calcium, and cell adhesion (**Figure 2B**). Regarding ontologies for molecular function (MF), proteins down-regulated in M2 were enriched for hyaluronic acid binding, while proteins up-regulated in M2 were enriched for actin monomer binding, copper ion binding, low-density lipoprotein binding, hexosaminidase activity, and fatty acid binding (**Figure 2C**). Both up and down-regulated proteins in M2 were enriched for biological processes related to the immune system (**Supplementary File 1**). Triglyceride catabolism, fibrinolysis, transmembrane ion transport, metabolism of reactive oxygen species, cation transmembrane transport, regulation of muscle system process, and peptidase activity are up-regulated in M2 (**Figure 2D**). In contrast, proteins associated with extracellular matrix (ECM) organization and cell adhesion were down-regulated in M2 group (**Figure 2D**). Cellular component (CC) analysis revealed that proteins up-regulated in M2 are enriched for the lysosome, cytoplasmic vesicles, and MHC complex. On the other hand, proteins down-regulated in M2 belong to the Golgi complex and membrane components (**Figure 2E**). Gene ontology protein network analysis showed that the ontology terms down-regulated in M2 form sub-networks related to immune system and cell adhesion processes, while ontologies up-regulated in M2 cluster into more diverse sub-networks, including lysosomal, secretory, iron homeostasis, triglyceride catabolism, and reactive oxygen metabolic-related processes (**Figure 2F**).

**Figure 2.**
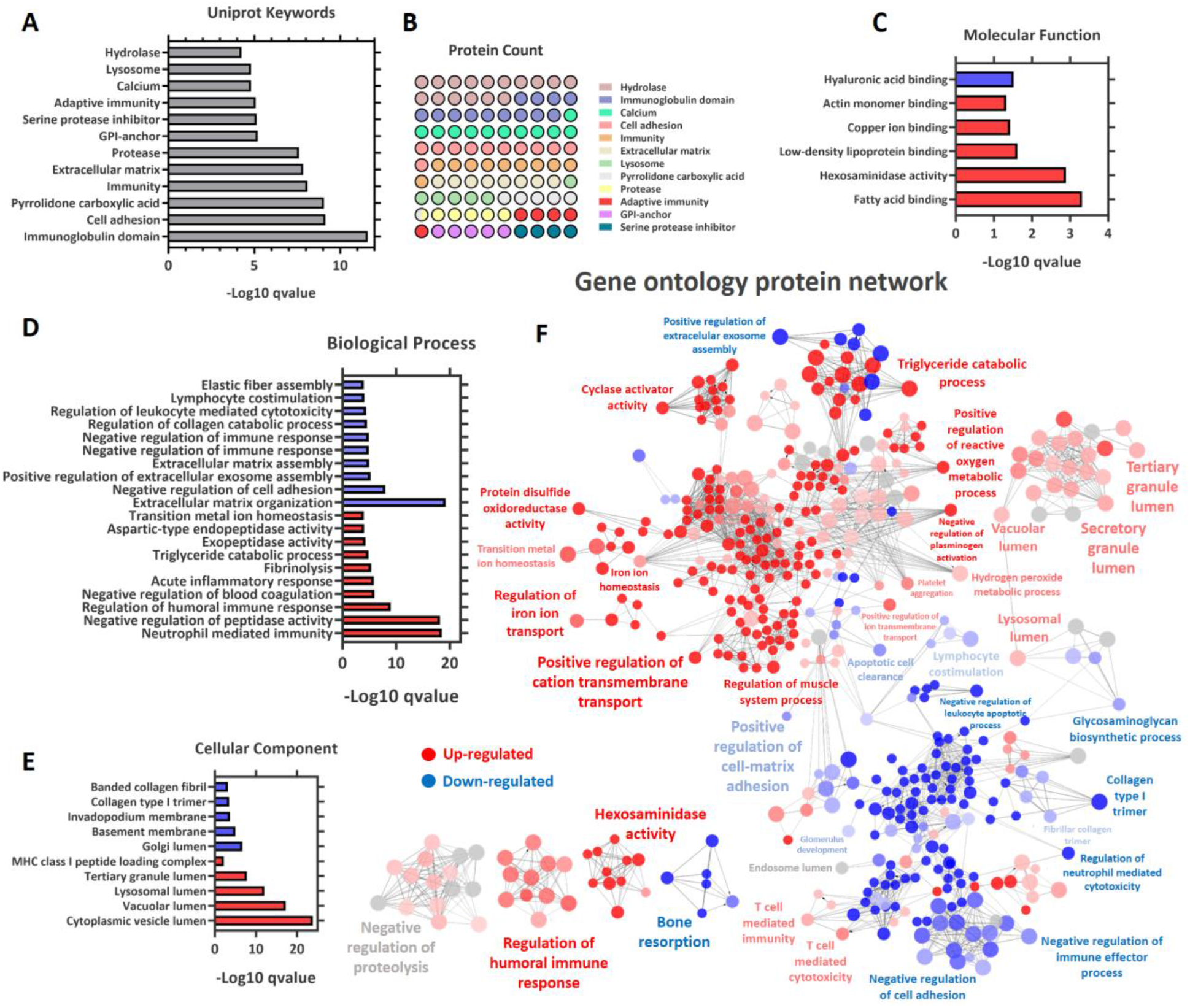
Urinary proteome remodeling during high physical stress reveals proteins related to different biological processes. UniProt keywords related to up-regulated and down-regulated proteins. Only terms with q-value <0.001 and protein count > 10 were considered (Benjamini-Hochberg) (**A**). Protein count for enriched keywords (**B**), GO Molecular function (**C**), GO biological process (**D**), GO cellular components (**E**), and interaction protein network (**F**) related to upregulated (red) and downregulated (blue) proteins. To determine ontologies, the ClueGo app was set to the significance threshold of 0.05 with the Benjamini-Hochberg correction.

Pathway enrichment analysis was also carried out. According to what has been observed for BP, proteins up-regulated in M2 are enriched for pathways linked to the immune system. Other enriched pathways in the group are related to diseases and interactions of the cell surface on the vascular wall (**Figure 3A**). The pathways identified with negative regulation in M2 include the organization of the ECM, ECM-proteoglycans, and interactions of the integrin cell surface (**Figure 3B and C**). Pathways related to catabolic processes and lysosome enzymes are enriched in M2 group (**Figure 3C**).

**Figure 3.**
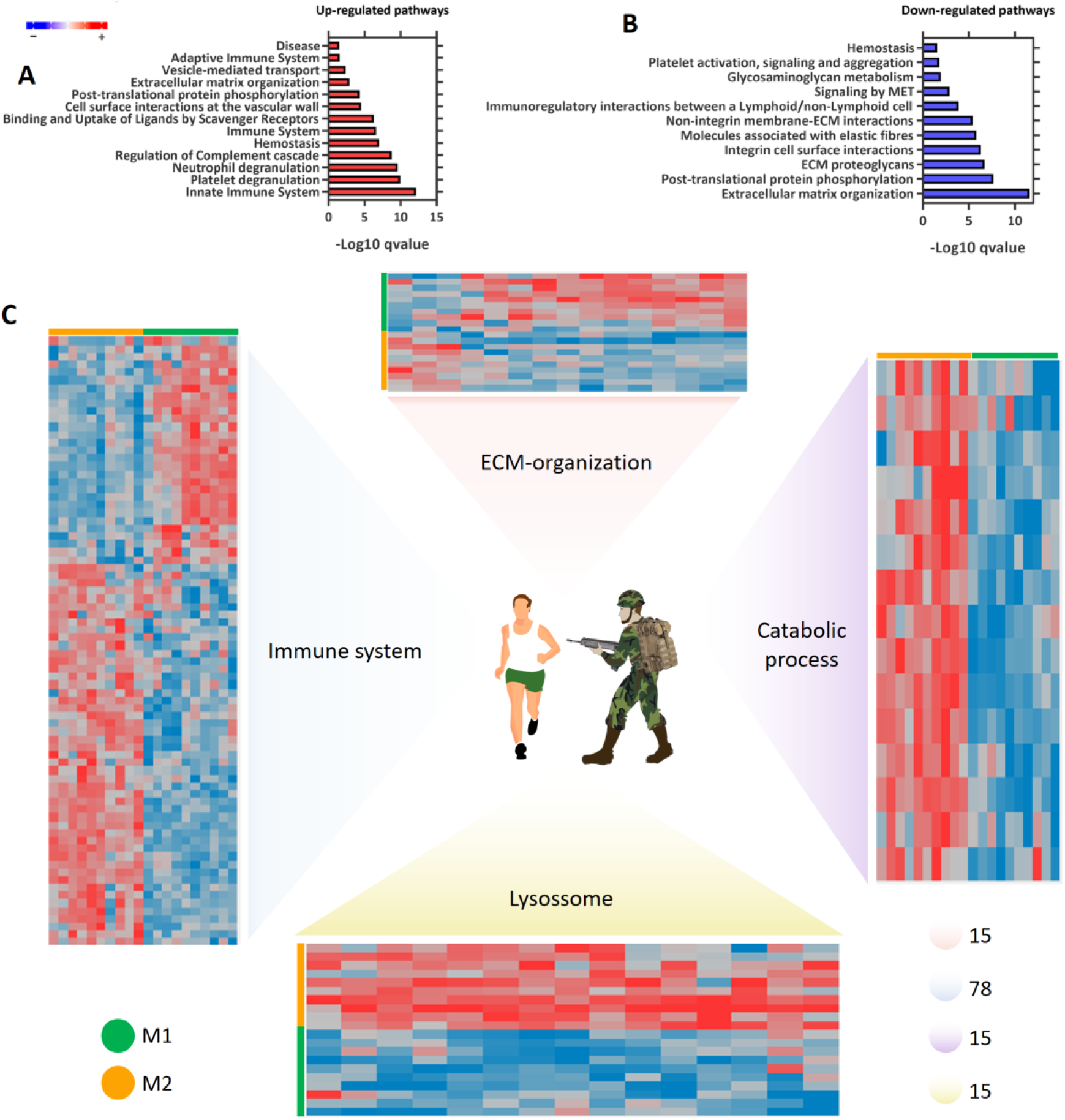
Pathway modulation related to extraneous exercise. Statistically enriched pathways related to positively regulated proteins (**A**) and negatively regulated proteins (**B**). The analyzes were performed with the g: profiler tool. The significance threshold chosen was 0.05 with the Benjamini-Hochberg correction. Heatmap showing the regulation of proteins related to immune system, extracellular matrix (ECM) organization, catabolic processes, and lysosome (**C**). The values were normalized using the z-score function. The red and blue colors show upregulated and downregulated proteins, respectively. The grey color indicates that the protein did not show any value in the sample. The number of differentially regulated proteins associated to each process are reported.

### Classic biochemical markers and differentially regulated proteins are correlated and indicate the progression of rhabdomyolysis (RM)

The classic biochemical markers of RM, MB, CK, AST, ALT, and LDH, show significantly (p-value < 0.01) higher levels in M2 when compared to M1 (**Figure 4**). Three of them (CK, LDH, and AST), widely used in the clinic to assess the progression of RM, were selected for correlation analysis to identify possible protein urinary biomarkers (**Table 1**).

**Figure 4.**
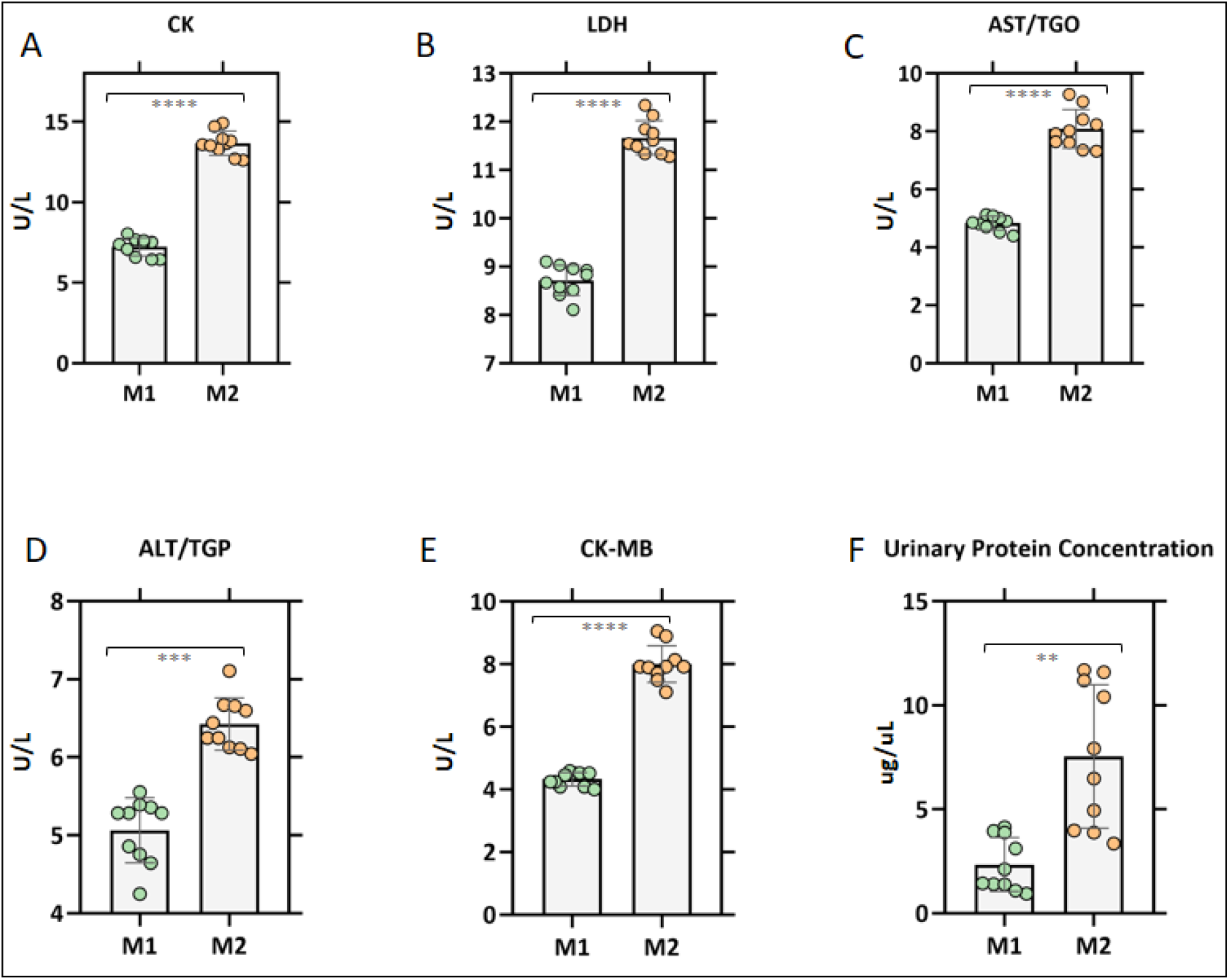
Biochemical parameters alteration in M1 and M2. Blood biochemical parameters associated to rhabdomyolysis are higher in participants during high physical stress mission M2 compared to M1. Classic rhabdomyolysis (RM) markers are indicated (**A-F**). All comparisons were statistically significant, with a p-value of <0.01.

**Table 1.** Correlation analysis between the three biochemical markers traditionally used for the diagnosis of rhabdomyolysis (RM) and urinary protein abundance.

|  | CK (U/L) |  | LDH (U/L) |  | AST (U/L) |  |
| --- | --- | --- | --- | --- | --- | --- |
|  | *r* | *p-value* | *r* | *p-value* | *r* | *p-value* |
|  | *spearman* |  | *spearman* |  | *spearman* |  |
| CTSH*** | 0,733 | 0,02023 | 0,648 | 0,04898 | 0,673 | 0,03897 |
| PIK3IP1 *** | -0,697 | 0,03056 | -0,782 | 0,01053 | -0,661 | 0,0438 |
| DEFB1*** | -0,673 | 0,03897 | -0,782 | 0,01053 | -0,661 | 0,0438 |
| ITGB1 *** | -0,661 | 0,0438 | -0,782 | 0,01053 | -0,721 | 0,02336 |
| BCAN *** | -0,661 | 0,0438 | -0,709 | 0,02675 | -0,709 | 0,02675 |
| TNFRSF10C *** | -0,661 | 0,0438 | -0,697 | 0,03056 | -0,648 | 0,04898 |
| GLRX1** | -0,709 | 0,02675 | -0,6 | 0,07343 | -0,77 | 0,01256 |
| BLVRB** | -0,685 | 0,03465 | -0,552 | 0,10488 | -0,685 | 0,03465 |
| DSC2** | -0,648 | 0,04898 | -0,588 | 0,08061 | -0,661 | 0,0438 |
| CO1A1** | -0,636 | 0,05443 | -0,673 | 0,03897 | -0,721 | 0,02336 |
| S28A1** | -0,636 | 0,05443 | -0,745 | 0,01741 | -0,648 | 0,04898 |
| GGCT* | 0,636 | 0,05443 | 0,685 | 0,03465 | 0,624 | 0,06032 |
| PGRP2** | 0,576 | 0,08828 | 0,661 | 0,0438 | 0,661 | 0,0438 |
| UTER** | -0,539 | 0,11385 | -0,673 | 0,03897 | -0,697 | 0,03056 |
| HBA** | 0,539 | 0,11385 | 0,648 | 0,04898 | 0,648 | 0,04898 |
| HMCN1* | -0,624 | 0,06032 | -0,733 | 0,02023 | -0,636 | 0,05443 |
| CCN3* | -0,6 | 0,07343 | -0,709 | 0,02675 | -0,612 | 0,06674 |
| PARK7* | 0,6 | 0,07343 | 0,685 | 0,03465 | 0,564 | 0,09628 |
| ICAM2* | -0,588 | 0,08061 | -0,661 | 0,0438 | -0,576 | 0,08828 |
| SLUR1* | -0,576 | 0,08828 | -0,648 | 0,04898 | -0,455 | 0,19124 |
| CATZ* | 0,564 | 0,09628 | 0,697 | 0,03056 | 0,624 | 0,06032 |
| FABP5* | 0,564 | 0,09628 | 0,697 | 0,03056 | 0,588 | 0,08061 |
| ATRN* | -0,539 | 0,11385 | -0,673 | 0,03897 | -0,491 | 0,15482 |
| PPIA* | 0,503 | 0,14403 | 0,661 | 0,0438 | 0,588 | 0,08061 |

A Spearman test was applied, and (\*), (\*\*), (\*\*\*) indicate proteins that have a correlation with one, two, or three biochemical rhabdomyolysis markers, respectively. The p value for each correlation is presented.

The correlation analysis of urinary protein abundance and blood levels of classic biochemical marker of RM (CK, LDH, and AST) indicated a panel of proteins that have a significant correlation with one, two, or three of those established biochemical markers for RM (**Table 1**). Most proteins have a significant correlation with at least one classical biochemical marker (GGCT, HMCN1, CCN3, PARK7, ICAM2, SLUR1, CATZ, FABP5, ATRN, and PPIA). A total of 8 proteins showed a significant correlation with at least two biochemical markers (GLRX1, BLVRB, DSC2, CO1A1, S28A1, PGRP2, UTER, and HBA). Proteins that have a significant correlation with three biochemical markers of RM (CTSH, PIK3IP1, DEFB1, ITGB1, BCAN, and TNFRSF10C) were selected as potential urine biomarkers. These proteins belong to different families and can play a role on the pathogenesis of RM by different mechanisms.

### Identification of a panel of urinary proteins capable of discriminating M1 and M2 with high specificity and sensitivity

The ROC curve of the proteins that correlate with the three selected classic biochemical marker of RM was evaluated (**Figure 5**). All proteins had sensitivity and specificity greater than 80%. The area under the curve (AUC) of all proteins was greater than 0.9, except for P3IP1 (**Supplementary File 1**). The PGCB and DEFB1 proteins had the highest AUC values (0.99). The PCA analysis based only on the protein panel shows that the quantitative proteome can separate the samples from M1 and M2 (**Figure 5H**).

**Figure 5.**
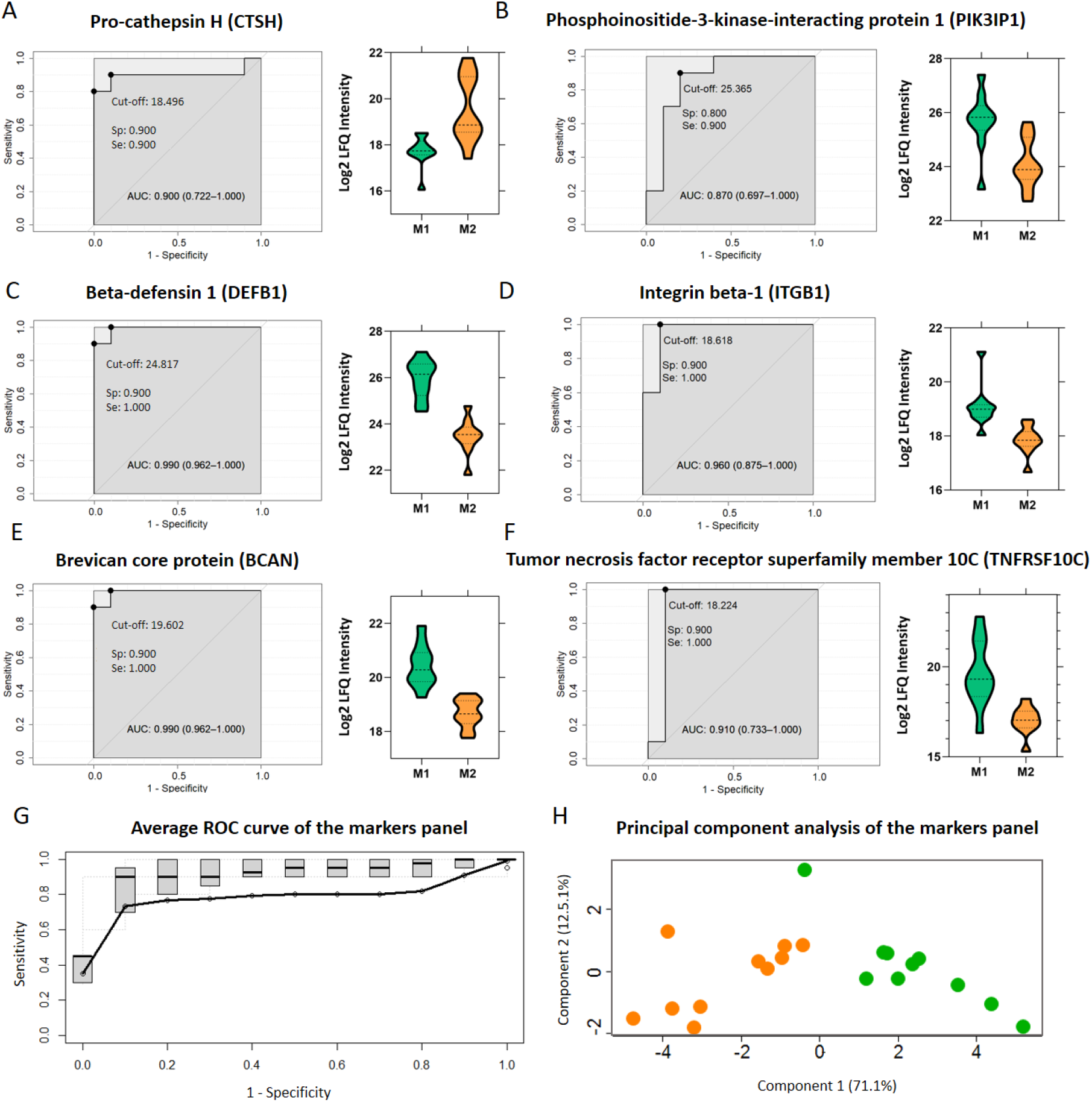
Evaluation of the performances of selected urinary protein markers to separate paritcipants subjected to high and low physical stress. ROC curves for a selected panel of rhabdomyolysis (RM) progression markers (**A-F**). Fusion of the individual ROC graphs (dotted in gray) of the marker proteins. The black line corresponds to the vertical average curve (**G**). Principal component analysis performed using only the selected panel of protein markers (**H**).

### Changes in the urinary proteome between genotypes modulating susceptibility to muscle damage

It is well known that certain genotypes are associated with protection or increased risk of muscle damage after physical exercise. The AGT MM and ACE II genotypes are associated with greater susceptibility to inflammation and muscle damage after physical exercise, and can be used to predict the development of muscle damage after strenuous exercise (Sierra et al., 2019). Therefore, we decided to determine the influence of *AGT M/T* and *ACE I/D* polymorphisms on the urinary proteome (**Figure 6**). The samples that were collected at M2 were subdivided into groups according to the referred genetic polymorphisms and changes in the proteome were assessed by T-test. The comparison between *AGT* MM (n = 3) and *AGT* TT (n = 5) genotypes showed a total of 15 differentially regulated proteins, with 7 upregulated and 8 downregulated (**Figure 6A**). Concerning *AGT* MM (n = 3) and *AGT* MT (n = 2) differences, a total of 20 regulated proteins were significantly regulated, of which 13 were up and 7 downregulated (**Figure 6B**). The comparison between the genotypes with less damage *AGT* MT (n = 2) and *AGT* TT (n = 5) genotypes revealed 2 upregulated and 9 downregulated proteins (**Figure 6C** and **Supplementary File 2**). CTSH, which has been previously identified as being able to differentiate M1 and M2 with high specificity and sensitivity (**Figure 5A-B**), was identified among the proteins with increased abundance in the genotypes with the highest risk for muscle damage (MM) (**Figure 6A and 6B**). The *ACE* I/D polymorphisms revealed 42 proteins regulated between *ACE* II (n = 2) and *ACE* DD (n = 5) genotypes, the majority being more abundant in the *ACE* DD genotype (32 proteins) (**Figure 6D**). Regarding *ACE* II (n = 2) and *ACE* ID (n = 3) genotypes, 29 proteins were differentially regulated, of which 5 and 24 are up- and down-regulated in the group with the highest susceptibility to damage (*ACE* II), respectively (**Figure 6E**). Comparison between genotypes associated to decreased risk for muscle damage (*ACE* ID and *ACE* DD) revealed 10 differentially regulated proteins (2 are up and 8 down-regulated) (**Figure 6F**). Taken together, these results show that the comparison of individuals with distinct genetic polymorphisms related to muscle damage susceptibility revealed several changes in the abundance of urinary proteins.

**Figure 6.**
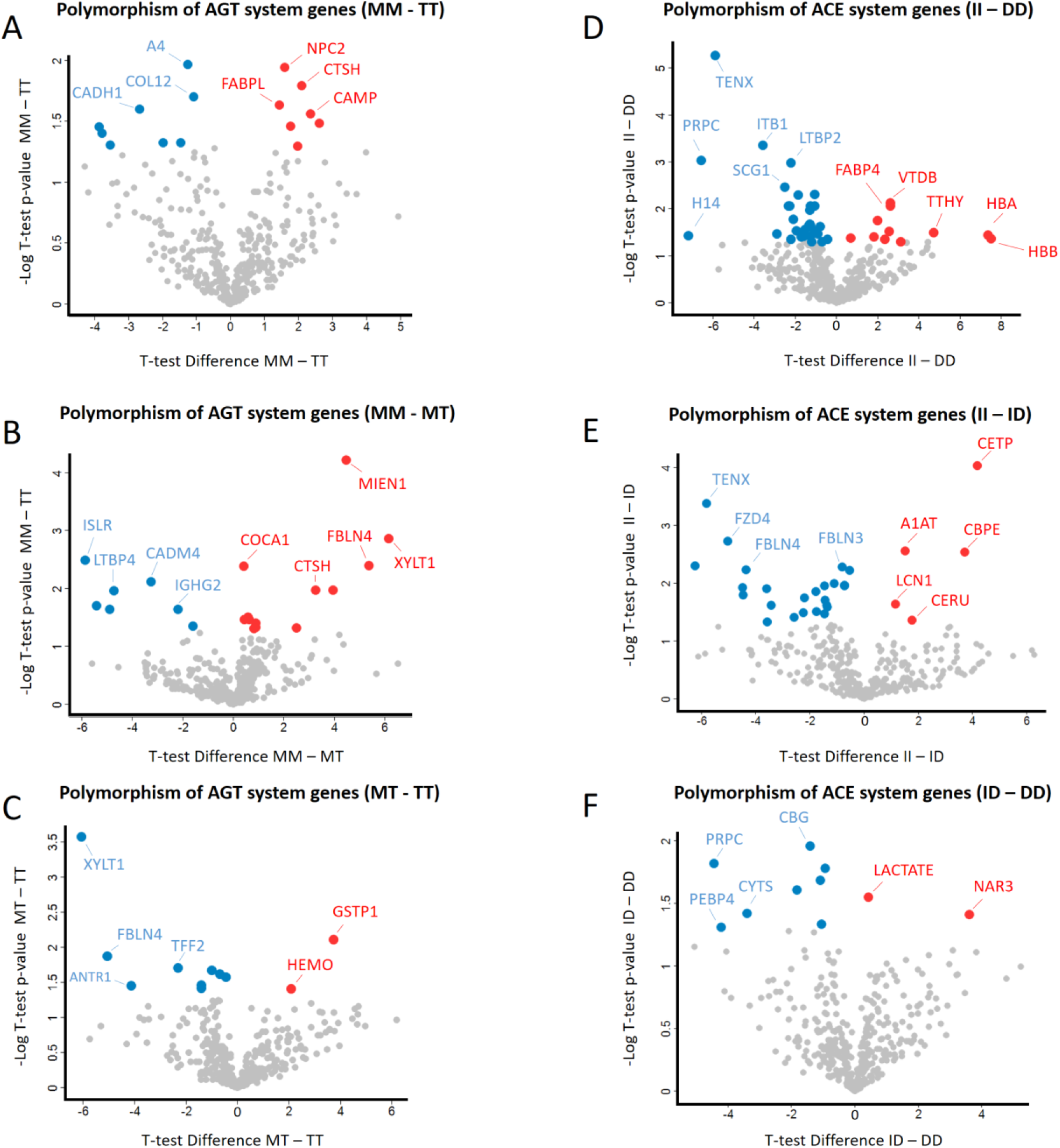
Changes in the urinary proteome between genotypes that modulate the susceptibility to rhabdomyolysis. Proteins differentially regulated between the polymorphisms: *AGT* (MM – MT – TT) (**A-C**) and *ACE* (II – ID – DD) (**D-F**). The differentially regulated proteins were determined by applying a student T-test, with the threshold of p-value ≤ 0.05. The red and blue colors show proteins with increased or decreased abundance, respectively.

Changes in the urinary proteome between individuals with high and low CK/inorganic phosphorus values was also performed (**Figure 7**). Samples that were collected at M2 were subdivided into groups according to CK values (CK < 10k and CK > 10k) and blood inorganic phosphorus concentration (P < 7 and P > 7) and T-test was applied. Thirty-three proteins and 5 biochemical variables are differentially regulated between individuals with CK > 10k and CK < 10k (**Figure 7A-B and Supplementary File 2**). Concerning the changes in biochemical variables, CK, LDH, AST/TGO, ACU, and FRAP are positively regulated in the CK > 10k group. The comparison based on blood phosphorus levels revealed that the group with values greater than 7 has 5 upregulated proteins, including OSTP (**Figure 7C-D**).

**Figure 7.**
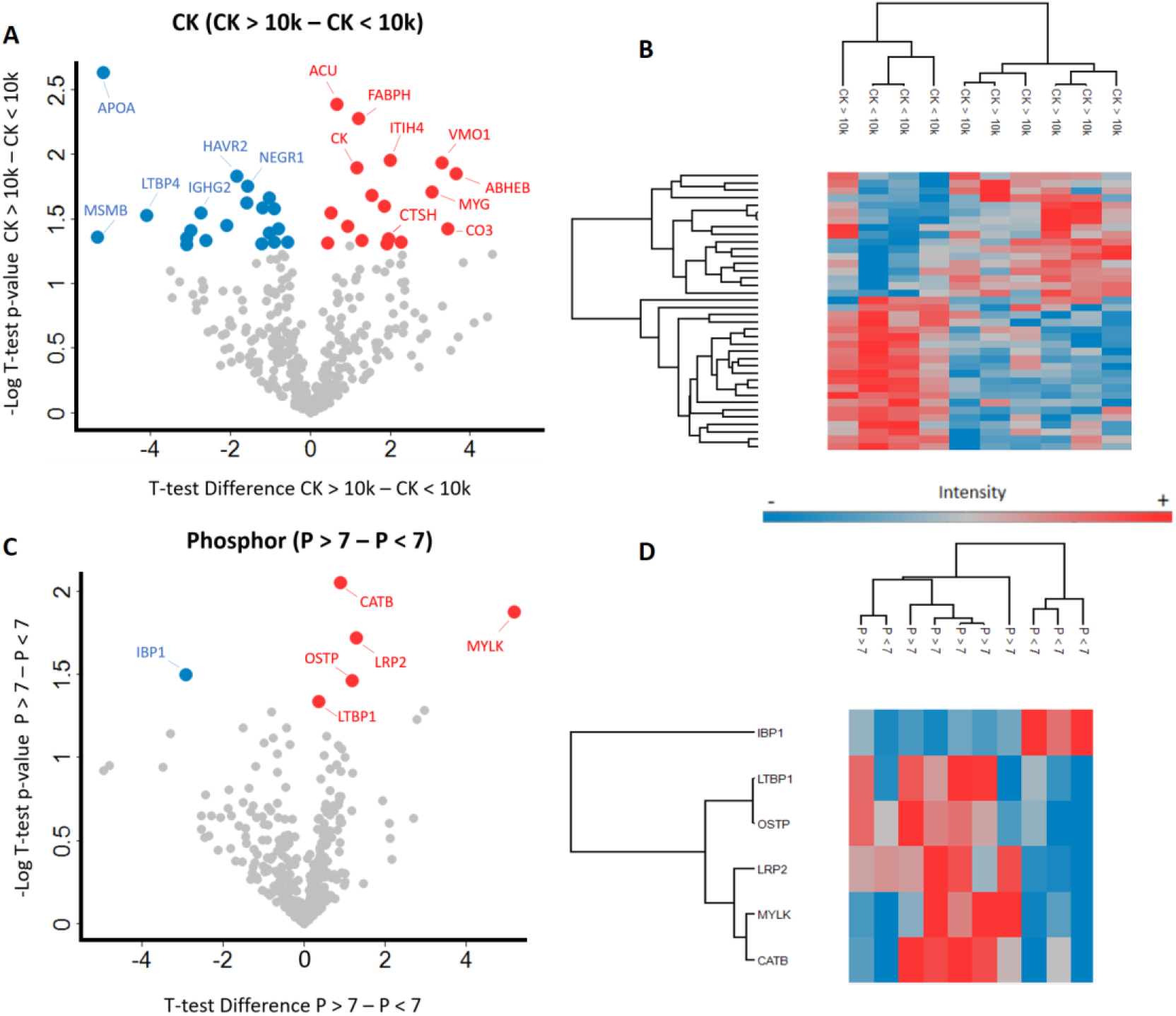
Changes in the urinary proteome between individuals with distinct levels of CK/inorganic phosphorus. Proteins differentially regulated between the parameters: CK (CK>10k - CK<10k) (**A-B**), and phosphorus (P >7 - P<7) (**C-D**). The differentially regulated proteins were determined by applying a paired student T-test, with the threshold of p-value ≤ 0.05. The red and blue colors represent positively and negatively regulated proteins, respectively.

The comparison of proteins that showed regulation based on genotypes and biochemical parameters shows that there were no differentially regulated proteins shared between CK and phosphorus (**Figure 8A**). On the other hand, there were 4 differentially regulated proteins in both genotype comparisons (*ACE* II genotype and *AGT* MM *genotype* polymorphism comparisons): PRDX6, PI16, VTNC, and FBLN4. Interestingly, all these proteins were regulated in the opposite direction between groups (**Figure 8B**). The comparison of upregulated proteins in groups with greater damage revealed that CTSH is common between comparisons of *AGT* MM *genotype* polymorphism and biochemical parameters. In addition, GC and SERPINA1 proteins were common between comparisons of *ACE* II genotype and biochemical parameters. Amongst the downregulated proteins in the groups susceptible to greater damage, 3 are shared between *AGT* MM *genotype* and biochemical comparisons (COLEC12, LTBP4, and IGHG2) and 6 between *ACE* II genotype and biochemical comparisons (BTN2A2, FBN1, NEGR1, NOTCH3, PIK3IP1, and PTGDS). It is worth mentioning that among this panel of differentially regulated proteins based on genotype and biochemical parameters, PIK3IP1 and CTSH, correlated with three of the most used blood biochemical markers for RM (**Table 1**) and had sensitivity and specificity greater than 80% to discriminate samples M1 and M2 (**Figure 5**), thus representing potential markers of muscle damage induced by physical exercise.

**Figure 8.**
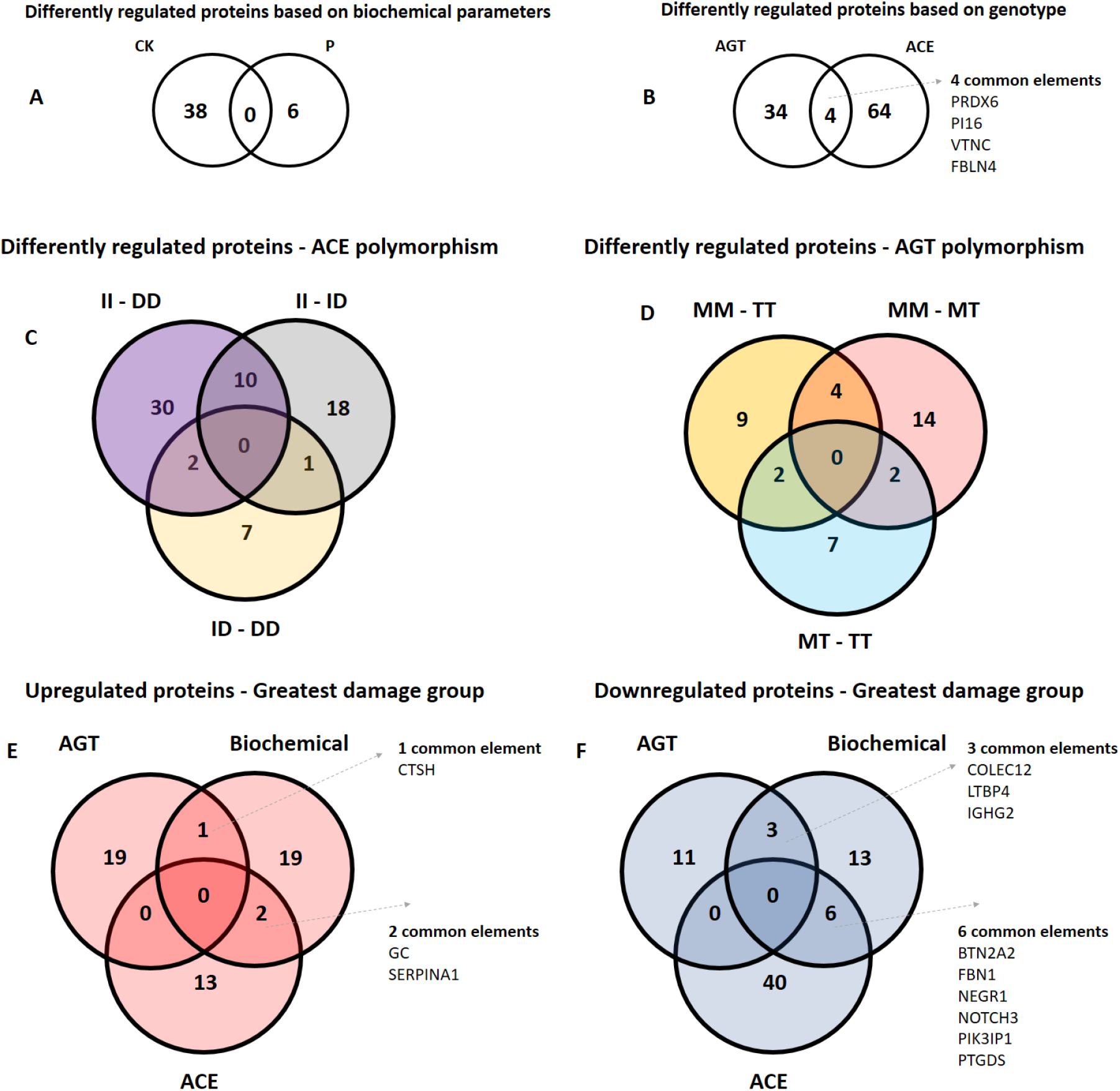
Comparison of proteins differentially regulated based on biochemical and genotypic parameters. Comparison of differentially regulated proteins based on biochemical variables (**A**) and genotype variables (**B-D**). Upregulated (**E**) and downregulated (**F**) proteins common in biochemical and genotypes groups with greater damage.

## Discussion

ERM is recurrent in military personnel due to the overload of military exercises and is associated with severe kidney and muscle damage (Torres et al., 2015). Currently, the diagnosis of RM is made through biochemical parameters measured in the blood (CK, MB, LDH, AST, and ALT). However, there is a great variation in the levels of these blood markers in individuals with RM, making an accurate diagnosis difficult. Therefore, identifying precise and accurate biomarkers in non-invasive biological matrices is essential to reduce the most alarming effects related to RM, which include renal failure and death (Petejova and Martinek, 2014). In this study, we performed a quantitative proteomic analysis of urine collected from military personnel during training in low and high strenuous physical exercise conditions. The identified upregulated proteins under intense physical exercise were mostly related to the immune system (CFB, F2, HPX, IGKC, IGKV3-15, IGKV3-20, IGKV4-1, IGLC3, IGLV1-47, IGLV3-25, PGC, SPINK5, VTN, C7, CD59), oxidative stress (PARK7, SOD1, GLRX, SH3BGRL3, TXN), and transmembrane cation transport (AGT, CALM3, F2, FLNA, GLRX, PARK7, TMSB4X).

During the severe effort, there is an increase in the production of cytokines and chemokines (Hamel et al., 2015). Tissue traumas associated with intense exercise have been associated with altered immunity (Lakier Smith, 2003). This contributes to severe infection process, and in some cases sepsis (Faist, 1996). In a study of a cohort of walkers participating in a walking event for 3 consecutive days, it was found that participants with activated neutrophils had an impaired physical sensation (Spijkerman et al., 2021). Interestingly, a case report showed that a 49 years old woman infected with HIV presented non-anion gap metabolic acidosis, acute kidney disease (AKI), RM, and hypokalemia indicating a hypergammaglobulinemic state (Bhargava et al., 2016).

Complement system activation associated with rhabdomyolysis-induced kidney injury was investigated in an animal model (Huang et al., 2018). The findings show that complement activation is associated with the onset of kidney damage, and that depletion of the complement system improves kidney and immune functions. This change in the complement system was confirmed in our study by the dysregulation of the CFAB and CO7 proteins, which act in the alternative and classical pathways, respectively. In M2, these proteins were downregulated, possibly due to their anti-inflammatory effects.

Studies have reported the presence of macrophages infiltrated in the kidneys of rats after induction of RM by glycerol and later confirmed in humans (Belliere et al., 2015). In addition, there is a close relationship between the immune system response and reactive oxygen species (ROS) production during physical exercise, since ROS can be produced due to immune response (Belliere et al., 2015; Simioni et al., 2018). It is known that regular exercise promotes immunity balance and has a positive outcome (Weyerer and Kupfer, 1994). However, under strenuous conditions, as in military training, rapid degeneration of skeletal muscle fibers and leakage of cellular content can occur, which is toxic to systemic circulation (Bagley et al., 2007).

The pathogenesis of RM includes an intense transport of intra and extracellular cations, characterized by increased levels of intracellular Ca^2+^. During physical exercise, skeletal muscle cells are excited and create an influx of Na^+^, resulting in similar amounts of K^+^ efflux. Influx and efflux activities strengthen the ability of Na^+^/K^+^-ATPase to adjust the concentrations of these cations within the cell; however, this ion distribution is dependent on ATP. If there is a decrease in available ATP, the levels of Na^+^ within the cell are increased due to a dysfunction of the Na^+^/K^+^-ATPase. Under conditions of equilibrium between Na^+^ and K^+^, the Na^+^/Ca^2+^ exchanger is activated and is responsible for the extrusion of Ca^2+^, but under conditions of increased intracellular Na^+^, the Na^+^/Ca^2+^ exchanger promotes the influx increasing the levels of Ca^2+^ within cell (Kim et al., 2016). The intracellular increase in Ca^2+^ is involved in the activation of proteases and phospholipases, which can act in the degradation of cell membrane phospholipids (Allen, 2004). In addition, this intracellular increase in Ca^2+^ is also associated with ROS generation, which can be toxic due to biomolecules oxidation (Cooper et al., 2002).

Other mechanisms that may be involved in the generation of ROS consist of the auto-oxidation of catecholamine hormones, which are released during exercise (Cooper et al., 2002). Ischemia-reperfusion mechanism is also associated with ROS and physical exercise. Physical activity is associated with hypoxia of different tissues, since blood is required in active skeletal muscles. In addition, during strenuous exercise, even muscle fibers may experience hypoxia. During tissue reoxygenation, ROS can be produced through the conversion of xanthine dehydrogenase in xanthine oxidase. Both enzymes act in the degradation of hypoxanthine to xanthine, however, only the action of xanthine oxidase results in the non-generation of ROs (Adiseshiah et al., 2005). Reactions related to the transformation of the heme group and the release of catalytic iron are also associated with the generation of ROS (Duvigneau et al., 2019). Here, we identified regulated proteins related to the detoxification of the heme group (BLVRD, FABPL, HPT, and LRP1), as well as proteins related to oxidative stress (UBA52, ATOX1, SOD1, TXN, CA1, and DDAH2).

Amongst the proteins that positively correlate with three classic biochemical marker of RM, we found cathepsin H (CTSH), which belongs to the superfamily of papain-like cysteine proteases and is involved in the degradation of intracellular proteins and ECM, as it is one of the main lysosomal enzymes (Chavez et al., 2016; Salminen et al., 1984). In addition the CTSB and CTSD proteins, also from the cathepsin family, were identified upregulated during high physical stress training. A previous study showed that cathepsin C (CTSC) expression was increased in rats submitted to strenuous exercise (Salminen et al., 1984). In addition, another study demonstrated that cathepsin K (CTSK) plays a key role in the loss of skeletal muscle and fibrosis, possibly through the reduction of inflammatory events (Hardy et al., 2016). To date, the increase in CTSH abundance caused by strenuous exercise has not yet been reported in the literature. However, we hypothesize that their functions related to degradation may be related to RM progression through cellular membrane degradation. Thus, our study reports for the first time the identification of increased levels of CTSH after strenuous exercise. A relationship has been described between lysosomal cathepsins and renal pathology, one of the most lethal complications of RM (Cocchiaro et al., 2017). The CTSB and CTSD proteins regulate ECM homeostasis, autophagy, apoptosis, glomerular permeability, endothelial function, and inflammation (Cocchiaro et al., 2017).

PIK3IP1 is a cell surface protein that is involved in the regulation of the PI3K pathway (Chen et al., 2019). Here, PIK3IP1 was downregulated after strenuous exercise, and this is the first study that correlates the abundance of this protein to muscle damage. Recently, PIK3IP1 was shown to be a regulator of inflammatory response of mesenchymal cells induced by TNFα (Brandstetter et al., 2019). In addition, it has been demonstrated that PIK3IP1 is an immune regulator that inhibits T cell responses (Chen et al., 2019), which is associated to the highest activity of T cells during high physical stress (Suzuki and Hayashida, 2021). PI3K is a highly conserved lipid kinase that is involved in physiological cardiac hypertrophy (PHH) (exercise training hypertrophy). Several signaling pathways have been proposed for different types of hypertrophy (Song et al., 2015). According to Song et al, 2015 PIK3IP1 deficiency was shown to lead to activation of the PI3K/protein kinase B (AKT)/mammalian target of rapamycin (mTOR) signaling pathway, increasing protein synthesis and cell size. According to this study, PIK3IP1 plays a negative compensatory role for PHH. PI3K is known to play a central role in regulating proliferation, cell growth, and cell survival. In particular, PI3K has been shown to regulate organ and cell size. Accordingly, we suggest that the downregulation of PIK3IP1 increased AKT activity and induced cardiomyocyte hypertrophy in volunteers through activation of the mTOR pathway, thus causing physiological hypertrophy in military personnel.

DEF1 belongs to the family of defensins, which are antimicrobial and cytotoxic peptides (Kagan et al., 1994). This family of peptides is associated with immune defense against bacteria, fungi, and viruses (Machado and Ottolini, 2015). Although the effects of inducing an immune response are well characterized, the involvement of the defensin family in anti-inflammatory activities have also been documented (Fruitwala et al., 2019). Here, we identified that DEF1 was downregulated in M2, which is associated with accumulation of muscle damage due to the strenuous physical exercise. It has already been reported the relationship between DEF1 and oxidative stress (Suzuki et al., 2011). In this context, the high DEF1 expression levels result in the increased on the production of glutathione, an important (tri)peptide that protects cells against oxidative stress. In this study we show that reduced downregulation of DEF1 could be associated with decreased production of GSH and, then with the impairment to protect muscle cells against oxidative stress generated by strenuous physical activity. Thus, our results indicate that oxidative stress seems to be highly relevant process in M2.

The ITGB1 protein is a member of the integrin family and is associated with cell adhesion. Here, this protein was identified downregulated in M2. This is in contrast with previous findings showing an increase in the abundance of ITGB1 after male Sprague-Dawley rats were subjected to a high load of physical exercise (Ogasawara et al., 2014). In addition, another study indicated that α7β1 integrin is related to the sensitization of skeletal muscle in response to mechanical tension and hypertrophy (Zou et al., 2011). Other studies have shown that α7β1 mRNA was positively regulated after eccentric contractions conferring protection to injuries from repeated exercise sessions (Boppart et al., 2008; Boppart and Mahmassani, 2019). Once the levels of α7β1 were decreased in M2, our data suggests that this protective effect has been lost after the strenuous exercise.

Brevican core protein (BCAN) is a neural proteoglycan that contributes to the formation of the ECM and is associated with several processes of physiological and pathophysiological plasticity in the brain (Frischknecht and Seidenbecher, 2012). According to the Human Protein Atlas database (Thul and Lindskog, 2018) BCAN mRNA is highly abundant in the brain, and it is also expressed in blood immune cells, including basophils. Here, we report for the first time a reduced abundance of BCAN protein in urine after strenuous physical activity. We hypothesize that this might reflect changes in the immune system due to physical exercise. Chondroitin sulfate proteoglycans (CSPGs) are the main components of ECM and have several functional roles. For example, CSPGs are generally secreted from cells and are structural components of a variety of human tissues, including cartilage, and are known to be involved in certain cellular processes, such as cell adhesion, cell growth, receptor binding, cell migration and interaction. with other ECM constituencies. (Murphy and Ohlendieck, 2016). Among the different types of macromolecules that participate in skeletal muscle biology, proteoglycans (PGs) have gained great attention in recent years and there are still many open questions about other roles during skeletal muscle physiology. Live expression during skeletal muscle development, repair, and maintenance reflects a multitude of specific activities. These functions can range from the modulation of cytokines, proteinases, cell adhesion molecules and growth factors. They participate in cell adhesion and migration or have recently emerged unexpected functions at the neuromuscular junction (JNM). At the cellular level, the location of PGs is critical, reflecting their different functions. Mikami et al., 2012 identified a signaling pathway responsive to the decline in CS abundance giving valuable insight into the role of CS as a critical extracellular signaling pathway that can be recognized by myogenic cell lines and regulates their differentiation. The PI3K/AKT pathway has been reported to be involved not only in vitro myogenesis but also in skeletal muscle regeneration. Notably, activation of AKT signaling has been shown to promote myofiber regeneration after CTX-induced injury and to neutralize MDX pathogenesis. Therefore, the regenerative effects of ChABC on skeletal muscle in vivo can also be exerted, at least in part, via the PI3K/AKT pathway. Further investigation is needed to understand the molecular basis of CS functions in cell-cell fusion and skeletal muscle regeneration processes (Mikami et al., 2012). Skeletal muscle formation and regeneration require the fusion of myoblasts to form multinucleated myotubes or myofibers, but their molecular regulation remains incompletely understood (Mikami et al., 2012). A study by Mikami et al., 2012 demonstrated that forced down-regulation of chondroitin sulfate chains increased myogenic differentiation *in vitro*. The study by Mikami et al., 2012, also demonstrated that the temporal decline of CS levels is an essential requirement for the processes of differentiation and repair of skeletal muscle (Mikami et al., 2012). Thus, we hypothesize that BCAN downregulation in M2 is related to muscle regeneration after strenuous physical activity.

TNFRSF10C is an apoptosis-inducing protein frequently studied in the context of cancer (Zhou et al., 2018). In addition, the tumor necrosis factor superfamily plays important roles in the activation, proliferation, differentiation and migration of immune cells to the central nervous system (CNS) (Vanamee and Faustman, 2018). It has been reported that TNFRSF10C mRNA of skeletal muscle biopsies from 25 men was increased after acute and long term exercise (Pourteymour et al., 2017). Accordingly, we detected reduced protein levels of TNFRSF10C in M2, but the role of the TNFRSF10C during physical exercise is still unknown. It was reported that TNFRSF10C mRNA from skeletal muscle biopsies from 25 men was increased after acute and long-term exercise (Pourteymour et al., 2017). TNFRSF10C is an antagonist decoy receptor and lacks the cytoplasmic death domain. TNFRSF10C protects from TRAIL-induced apoptosis by competing with DR4 and DR5 for binding to TRAIL (Schmoldt et al., 1975). TRAIL is a TNF superfamily cytokine recognized for its ability to trigger selective apoptosis in tumor cells, although it is relatively safe compared to normal cells. It’s binding to its cognate agonist receptors, namely death receptor 4 (DR4) and/or DR5, can induce the formation of a membrane-bound macromolecular complex, coined DISC (death signaling induction complex), necessary and sufficient for involving the apoptotic machinery. TNFRSF10C appears to be a p53 target gene that is regulated by genotoxic stress in a p53-dependent manner. Thus, reduced levels of TNFRSF10C in M2 may indicate that the apoptotic process is dysregulated (Micheau, 2018), (Sheikh et al., 1999; Sheridan et al., 1997).

The fatty-acid binding proteins family stood out in the analysis of the urinary proteome after strenuous exercise (FABP1, FABP3, FABP4, FABP5). These proteins are involved in the binding and transport of fatty acids into specific cellular organelles for lipid storage and metabolism and catabolic processes. Several fatty-acid binding proteins were up-regulated in M2. Amongst the identified fatty-acid binding proteins, the FABP3 (also known as FABPH) presented the highest abundance in urine after strenuous exercise. It has been reported that Endurance training increases the expression of fatty acid binding protein in human skeletal muscle and this is associates to increased metabolism of fatty acids and carbohydrates during exercise (Higashi et al., 1997; Tunstall et al., 2002). Several lines of evidence have shown that serum FABP3 is a valuable blood biomarker for detection of muscle lesions, modulating the levels of fatty acids in the cells of skeletal muscle, brain, liver, intestine and heart (Burch et al., 2015, Pritt et al., 2008, Burch et al., 2016, Zhang et al., 2017, Goldstein, 2017, Fiuza-Luces et al., 2018). Another study showed that serum FABP3, together with other skeletal muscle serum biomarkers, was predictive for necrosis of skeletal muscle (Pritt et al., 2008). Accordingly, when serum FABP3 was used in conjunction with classical biomarkers of muscle lesion, such as CK and AST, it improved the sensibility and specificity of the diagnostics of skeletal muscle lesion induced by drugs in rats (Burch et al., 2016). FABP3 had a higher association with muscle weakness in polymyositis and dermatomyositis patients than CK and myoglobin serum levels (Zhang et al., 2016). A recent study showed that serum levels of FABP3 might represent a novel biomarker for Duchenne muscular dystrophy (Dowling et al., 2020). To the best of our knowledge, this is the first report that shows an increased abundance of FABP3 in human urine after strenuous exercise and its correlation with serum biomarkers of muscle damage. In addition, the abundance of FABP5 in urine showed a positive correlation with LDH blood level, a classic RM marker.

Continuously delivering ATP to the fundamental cellular processes that support skeletal muscle contraction during exercise is essential for sporting performance in events lasting from seconds to several hours. Because muscle ATP stores are small, metabolic pathways must be activated to maintain the required rates of ATP re-synthesis. These pathways include the degradation of phosphocreatine and muscle glycogen, thus allowing for substrate-level phosphorylation (‘anaerobic’) and oxidative phosphorylation using reducing equivalents of carbohydrate and fat metabolism (‘aerobic’) (Hargreaves and Spriet, 2020). ATP is required for the activity of the main enzymes involved in membrane excitability (Na + / K + ATPase), sarcoplasmic reticulum calcium manipulation (Ca 2+ ATPase), and myofilament cross-bridge cycling (myosin ATPase). As intramuscular ATP stores are relatively small (~5 mmol per kg of wet muscle), they are unable to sustain contractile activity for long periods (Hargreaves and Spriet, 2020). Therefore, other metabolic pathways must be activated, including substrate-level (or anaerobic) phosphorylation and oxidative (or aerobic) phosphorylation. Anaerobic energy pathways have a much higher potency (ATP production rate) but a lower capacity (total ATP produced) than aerobic pathways (Sahlin et al., 1998). During events that last for several minutes to hours, the oxidative metabolism of carbohydrates and fats provides almost all of the ATP for skeletal muscle contraction. Even during marathon and triathlon events lasting 2–2.5 h, there is a primary dependence on carbohydrate oxidation (Hawley and Leckey, 2015). The main intramuscular and extramuscular substrates are muscle glycogen, blood glucose (derived from hepatic glycogenolysis and gluconeogenesis and intestine when carbohydrate is ingested), and fatty acids derived from both muscles (intramuscular triglyceride (IMTG)) and tissue triglyceride reserves adipose (Hargreaves and Spriet, 2020). An increase in adipose tissue lipolysis supports the progressive increase in plasma fatty acid uptake and oxidation (Horowitz and Klein, 2000), but as lipolysis exceeds uptake and oxidation, plasma fatty acid levels increase. Inhibition of adipose tissue lipolysis increases dependence on both muscle glycogen and IMTG but has little effect on muscle glucose uptake (van Loon et al., 2005). The importance of IMTG oxidation during exercise has been a matter of debate, and results in the literature may be influenced by differences in individuals’ training status, sex, fiber type distribution, and resting IMTG stores (Kiens, 2006). However, IMTG appears to be an important source of fuel during exercise in trained individuals. Thus, we hypothesize that the increase in FABPs in M2 is due to the intense lipid catabolism that occurs and is necessary for the generation of energy for myocytes during strenuous physical activity.

Elucidation of the mechanisms involved in muscle damage and inflammation during strenuous physical exercise is important to predict clinical complications. Recently, it has been reported that polymorphisms of the angiotensin-converting related enzyme impact on inflammation muscle and myocardial damage during endurance exercise (Sierra et al., 2019). We show herein the association of polymorphisms of the angiotensin-converting enzyme (ACE) I/D and angiotensinogen (AGT) Met235Thr with the urinary proteome after strenuous physical exercise.

Metabolic stress can influence muscle damage and exercise-induced inflammatory responses. The polymorphisms of the angiotensin-converting enzyme (ACE) I/D gene showed an increase in the abundance of proteins related to oxidative stress (HBB, PRDX6, CALM3, PPIA and FABP4) compared to the *ACE* II genotype, being the latter correlated with higher risk to develop muscle damage (Yamin et al., 2007). The ACE DD and AGT TT genotypes were associated with low muscle damage and indicate that the renin-angiotensin-aldosterone system (RAS) plays an important role in exercise-induced muscle damage (Sierra et al., 2019). The RAS is responsible for regulating important processes, which include blood pressure control, water, electrolyte balance and recent evidence suggests impacts on skeletal muscle wasting (Dai et al., 2019). However, this important physiological system is also related to pathological processes, such as inflammation and oxidative stress (Ramalingam et al., 2017). It is proposed that angiotensin II is a potent stimulator of ROS production, which results in the oxidation of low-density lipoprotein (LDL) and, therefore, directs the inflammatory effects of angiotensin II (Husain, 2015). In addition, there is a hypothesis that the increase in ROS activate the degradation of muscle proteins mediated by the proteasome system, which is activated and contributes to the atrophy of the skeletal muscle (Sukhanov et al., 2011). Oxidative stress mediated by the renin-angiotensin system has been shown to lead to mitochondrial dysfunction in different tissues, including heart, liver, kidney, muscle, adipose tissue and pancreas (Ramalingam et al., 2017). PRDX-6, an oxidoreductase ubiquitously expressed in the cytosol, detoxifies ROS and is a marker of oxidative damage. PRDX-6 was previously suggested to be an important transducer of H_2_O_2_ levels induced by exercise (Wadley et al., 2016). A recent work reported that among a panel of blood biomarkers associated with brain lesions, PRDX-6 presented the highest increase in abundance after high intensity interval training (HIIT) (Di Battista et al., 2018). The CTSH, CAMP and CYTB proteins were upregulated in participants genetically susceptible to greater damage due to the polymorphism of the *AGT* Met235Thr gene. CTSH correlated with three classic biochemical markers of rhabdomyolysis, being upregulated in the groups with the highest damage (genotype *AGT* MM and CK> 10K) or who practice strenuous exercise (M2), suggesting that there is higher degradation of the ECM and possible disruption due to strenuous physical exercise.

**Figure 9** is a synopsis of the main proteins found in urine after strenuous physical activity (M8). Proteins written in red are up while those in blue are down-regulated and in black the unregulated proteins. It is noteworthy that several cathepsins (CTSH, CTSD, CTSB and CTSZ) responsible for cell lysis and degradation of the extracellular matrix were identified as overexpressed, in addition to several proteins related to muscle fibers, including titin (TTN), one of the largest muscle fiber proteins, which is currently considered a biomarker of muscle injury (Kanda et al., 2017; Vidak et al., 2019; Laurens et al., 2020). We also identified the presence of several myokines (cathepsins) and muscle proteins (such as actin, myosins and philamine) including biomarkers of muscle injury (FABPH, MYO and TTN) (**Figure 9A**) (Kanda et al., 2017; Vidak et al., 2019; Laurens et al., 2020). It is worth mentioning the finding of overexpressed renal lesion biomarkers: (i) of podocytes and glomeruli (ORM1 and B2M); (ii) proximal tubule (LRG1e FABP1); and (iii) distal tubule (FABPL), although clinical and laboratory results have not identified kidney damage through creatinine) (Colhoun and Marcovecchio, 2018; Weber et al., 2017). In addition, many proteins related to the metabolism and catabolism of the heme were also differentially expressed. According to Johan Fevery’s study (2008), hemolysis is found from increased LDH and decreased free haptoglobin (down-regulated), as found in our results (**Figure 9**) (Fevery, 2008). Our hypothesis for this finding is that lysis of red blood cells and myocytes occurs during strenuous physical activity, leading to degradation of free hemoglobin and myoglobin in the cytosol, so that the heme group, product of this degradation, binds to hemopexin (up regulated). In this scenario, the heme group not connected to hemopexin is degraded by hemoxigenase-1 (not identified/not shown) to biliverdin, carbon monoxide (CO) and Fe^2+^. Biliverdin is degraded to bilirubin (antioxidant protein) by biliverdin reductase (BLVRB, up regulated), while carbon monoxide (CO) is degraded to carbon dioxide (CO_2_) and water (H_2_O) by carbonic anhydrase (CA1, up regulated). Hepcidin (overexpressed) is responsible for increasing the expression of ferroportin to prevent the exit of Fe^2+^. The extravasated Fe^2+^ participates in the fenton reaction, forming ROS. Several proteins related to the inflammatory process and oxidative stress are overexpressed after strenuous physical exercise, possibly to detoxify this insult (Carneiro et al., 2021; Van Avondt et al., 2019). Finally, we highlight the finding of proteins related to the activation of RAAS, such as angiotensinogen. This hormone system is characteristically induced by strenuous physical activity which is inherent to the special operations course that aims to train the elite troop of the Brazilian Navy.

**Figure 9.**
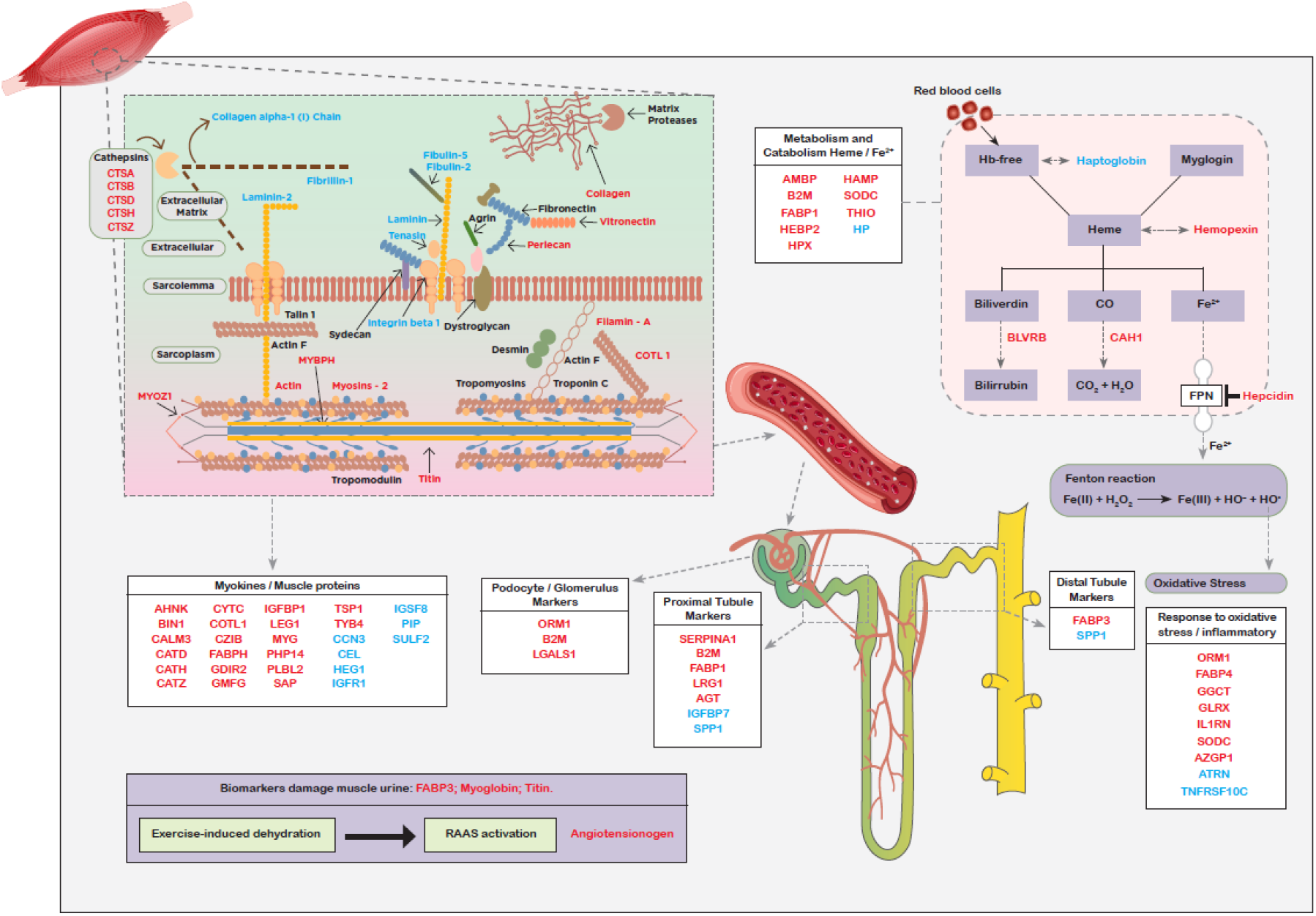
Synopsis of the urinary proteomic profile found in the analyzed missions. Sarcolemma proteins and sarcomere structure (A). Proteins and processes involved: Cathepsins causing ECM lysis, with protein leakage from myocytes into the blood stream (B). Biomarkers of muscle injury, myokines and muscle proteins (C). Biomarkers of lesions in podocytes and glomeruli (D), proximal (E) and distal tubules (F). Proteins related to heme metabolism and catabolism (G), oxidative stress and inflammatory process (H). Activation of the RAAS (angiotensinogen) system (I). Red: up-regulated proteins; Blue: down-regulated proteins; Black: unregulated proteins found in the urine of volunteers. Abbreviations: Hb, hemoglobin; CO, carbon monoxide; Fe^2+^, ferrous iron; CO, carbon monoxide, CO2, carbon dioxide; H_2_0, water; FPN, ferroportin; Fe(II), iron oxide; H_2_O_2_, hydrogen peroxide; Fe(III), ferric iron; HO^-^, hydroxide; HO^+^, hydroxylic cation; RAAS, renin-angiotensin-aldosterone system.

## Conclusions

The urinary proteome is an important opportunity to monitor pathophysiological changes in a non-invasive manner. In this study, we showed that the urinary proteome of military personnel undergoing strenuous training may provide a better understanding about the pathogenic mechanisms of rhabdomyolysis and its potential biomarkers. This is the first report that integrates genetic, biochemical, and urinary proteomic data of military subjected to intense physical training, providing a comprehensive overview of the disorders triggered by rhabdomyolysis. We identified potential protein biomarkers for rhabdomyolysis in the urine. Amongst these proteins, CTSH, involved in the degradation of ECM, has an excellent potential to discriminate groups exposed to higher muscle damage mediated by exercise. CTSH also distinguishes individuals that have higher genetic and biochemical susceptibility to muscle damage, a feature that other classical blood biochemical markers of rhabdomyolysis (CK, LDH, and AST) do not have. Strikingly, CTSH protein levels in the urine correlated with these three markers currently used in the detection of rhabdomyolysis. Other findings indicate that the greater overload of physical exercises affects the immune system and increases oxidative stress and ion transport in the transmembrane, processes that correlate with the progression of rhabdomyolysis. Taken together, our findings show the potential use of certain urine proteins as molecular markers of muscle damage due to intense physical conditions such as military training activities.

## Acknowledgements

The authors would like to thank to Brazilian Navy - *Marinha do Brasil (Comando do Material do Corpo de Fuzileiros Navais, Hospital Naval Marcilio Dias, Instituto de Pesquisas Biomédicas da Marinha*, *Centro de Educação Física Almirante Adalberto Nunes*, *Centro de Instrução Almirante Sylvio de Camargo* and *Centro de Tecnologia do Corpo dos Fuzileiros Navais*) for the research support and providing the facilities; We also thank to *Greiner Bio-One Brasil Produtos Médicos Hospitalares* and *Anil Lab 1288 Comércio e Representações* for provide the materials and reagents for biochemical analysis; *Coordenação de Aperfeiçoamento de Pessoal de Nível Superior* (CAPES), *Conselho Nacional de Desenvolvimento Científico e Tecnológico* (CNPq), *Fundação Carlos Chagas de Amparo à Pesquisa do Rio de Janeiro* (FAPERJ) and *Fundação de Amparo à Pesquisa do Estado de São Paulo* (FAPESP - 2018/18257-1, 2018/15549-1, 2020/04923-0 and 2014/27198-8) for providing financial support for this study.

## Conflict of interest

Andréia Carneiro da Silva, Marcos Dias Pereira and Giuseppe Palmisano declare a conflict of interest due to the filing of a patent in Brazil, part of the data in this article.

## Disclaimer

The views expressed are those of the authors and do not reflect the official policies of the Brazilian Navy, the Department of Defense, or the Brazil Government.

## Data available

The mass spectrometry proteomics data have been deposited to the ProteomeXchange Consortium via the PRIDE partner repository with identifier PXD030207.

## Supplementary Files

**Supplementary File 1.** Proteins identified by MaxQuant v.1.5.5.135 in urine samples (A); regulated proteins between groups M1 and M2. Description: protein description from the Swiss-Prot database. SwissProt / UniProt database name identifier (B); Enrichment analysis performed using DAVID, ClueGO, and g: profile (C-G) tools. The adjusted p-value refers to the application of the correction by the Benjamini-Hochberg method. ROC curve for ITGB1, CTSH, DEFB1, PIK3IP1, BCAN, and TNFRSF10C. AUC: Area under the curve (H).

**Supplementary File 2.** Proteins regulated between the genotypes II - DD (B), II - ID (C), ID - DD (D), MM - TT (E), MM - MT (F), and MT - TT (G). Description: protein description from the Swiss-Prot database. SwissProt / UniProt database name identifier. Venn diagram for AGT (H), ACE (I), Downregulated in the most damaging groups (J) Upregulated in the most damaging groups (L).

## Notes

### Competing Interest Statement

Andreia Carneiro da Silva, Marcos Dias Pereira and Giuseppe Palmisano declare a conflict of interest due to the filing of a patent in Brazil, part of the data in this article.

